# A single dose, BCG-adjuvanted COVID-19 vaccine provides sterilizing immunity against SARS-CoV-2 infection in mice

**DOI:** 10.1101/2020.12.10.419044

**Authors:** Claudio Counoupas, Matt D. Johansen, Alberto O. Stella, Duc H. Nguyen, Angela L. Ferguson, Anupriya Aggarwal, Nayan D. Bhattacharyya, Alice Grey, Karishma Patel, Rezwan Siddiquee, Erica L. Stewart, Carl G. Feng, Nicole G. Hansbro, Umaimainthan Palendira, Megan C. Steain, Bernadette M. Saunders, Jason K. K. Low, Joel P. Mackay, Anthony D. Kelleher, Warwick J. Britton, Stuart G Turville, Philip M. Hansbro, James A. Triccas

## Abstract

Global control of COVID-19 requires broadly accessible vaccines that are effective against SARS-CoV-2 variants. In this report, we exploit the immunostimulatory properties of bacille Calmette-Guérin (BCG), the existing tuberculosis vaccine, to deliver a vaccination regimen with potent SARS-CoV-2-specific protective immunity. Combination of BCG with a stabilized, trimeric form of SARS-CoV-2 spike antigen promoted rapid development of virus-specific IgG antibodies in the blood of vaccinated mice, that was further augmented by the addition of alum. This vaccine formulation, BCG:CoVac, induced high-titre SARS-CoV-2 neutralizing antibodies (NAbs) and Th1-biased cytokine release by vaccine-specific T cells, which correlated with the early emergence of T follicular helper cells in local lymph nodes and heightened levels of antigen-specific plasma B cells after vaccination. Vaccination of K18-hACE2 mice with a single dose of BCG:CoVac almost completely abrogated disease after SARS-CoV-2 challenge, with minimal inflammation and no detectable virus in the lungs of infected animals. Boosting BCG:CoVac-primed mice with a heterologous vaccine further increased SARS-CoV-2-specific antibody responses, which effectively neutralized B.1.1.7 and B.1.351 SARS-CoV-2 variants of concern. These findings demonstrate the potential for BCG-based vaccination to protect against major SARS-CoV-2 variants circulating globally.

## INTRODUCTION

The world has entered a critical stage in the continuing fight against COVD-19. The deployment of effective vaccines has had a profound impact in reducing cases and SARS-CoV-2 transmission in countries with high vaccine coverage^1,2^. However global cases are again on the rise, driven predominantly by a surge of infections in Europe, South America and the subcontinent due to the emergence of new SARS-CoV-2 variants. Two of the dominant variants of concern (VOCs), B.1.1.7 and B.1.351, both have increased transmissibility^3,4^ and antibodies from convalescent patients and vaccinees have a reduced capacity to neutralize B.1.351^5,6^. Vaccination with one of the most widely deployed vaccines, ChAdOx1 nCoV-19, does not protect against mild-to-moderate COVID-19 due to the B.1.351 variant in South Africa^7^. A critical issue is ensuring the adequate supply of vaccines to the hardest-hit low and middle income countries, particularly due to complex logistical requirements (e.g. storage at low temperature for mRNA vaccines). The requirement for multiple doses for most approved vaccines is a barrier to rapid, mass vaccination and has necessitated changes in dosing schedules in some countries to ensure sufficient vaccine coverage^8^. Thus, ensuring the global supply of vaccines effective against emerging variants will be necessary to control the global COVID-19 pandemic.

One unique strategy is to ‘repurpose’ existing licensed vaccines for use against COVID-19. Significant interest has focussed on *Mycobacterium bovis* bacille Calmette-Guerin (BCG), the tuberculosis (TB) vaccine. Considerable data has been accumulated to show that BCG has beneficial, non-specific effects on immunity that affords protection against other pathogens, particularly respiratory infections^9^. Most recently, BCG vaccination was shown to protect against viral respiratory tract infections in the elderly (greater than 65 years old) with no significant adverse events^10^. This non-specific protective effect is attributed to the ability of BCG to induce ‘trained immunity’ i.e. reprogramming of innate immune responses to provide heterologous protection against disease. For these reasons, a Phase 3, randomised controlled trial in healthcare workers has commenced to determine if BCG vaccination can reduce the incidence and severity of COVID-19 (the BRACE Trial)^9^. While that trial will determine if BCG can reduce the impact on COVID-19 during the current pandemic, BCG does not express SARS-CoV-2 specific antigens and thus, would not induce long-term immune memory.

Here, we have exploited the immunostimulatory properties of BCG to develop a SARS-CoV-2 vaccine, BCG:CoVac, that combines a stabilized, trimeric form of the spike protein with the alum adjuvant. BCG:CoVac stimulated SARS-CoV-2-specific antibody and T cell responses in mice after a single vaccination, including the elicitation of high-titre NAbs. Critically, a single dose was shown to protect mice against severe SARS-CoV-2, demonstrating that BCG:CoVac is a highly immunogenic and promising vaccine candidate.

## RESULTS

### BCG vaccination promotes SARS-CoV-2 specific antibody and T cell responses in mice

The immunostimulatory properties of BCG^11^ led to us to test if the vaccine could serve as the backbone for a unique vaccine platform against COVID-19. This was also supported by our observation that prior BCG immunization could augment anti-spike IgG responses after boosting with SpK formulated in Alhydrogel/alum (Alm^SpK^) (Fig. S1). To determine if this property of BCG could be used in a single vaccine formulation, we subcutaneously (s.c) vaccinated mice with a single dose of BCG formulated with a stabilized, trimeric form of the SARS-CoV-2 spike protein^12^ and the titre of IgG2c or IgG1 anti-SpK antibodies was determined at various timepoints post-immunization (Fig. 1a). While BCG vaccination resulted in background levels of anti-SpK antibodies, titres were approximately 100-fold higher for both antibody isotypes after BCG^Spk^ vaccination, and similar to those levels achieved with Alm^SpK^ (Fig. 1b, 1c). Addition of alum to BCG^Spk^ (termed BCG:CoVac) further increased antibodies titres, particularly IgG2c, which were significantly greater after BCG:CoVac vaccination compared to mice immunized with either BCG or Alm^SpK^, at all timepoints examined (Fig. 1b, 1c).

**Fig. 1.**
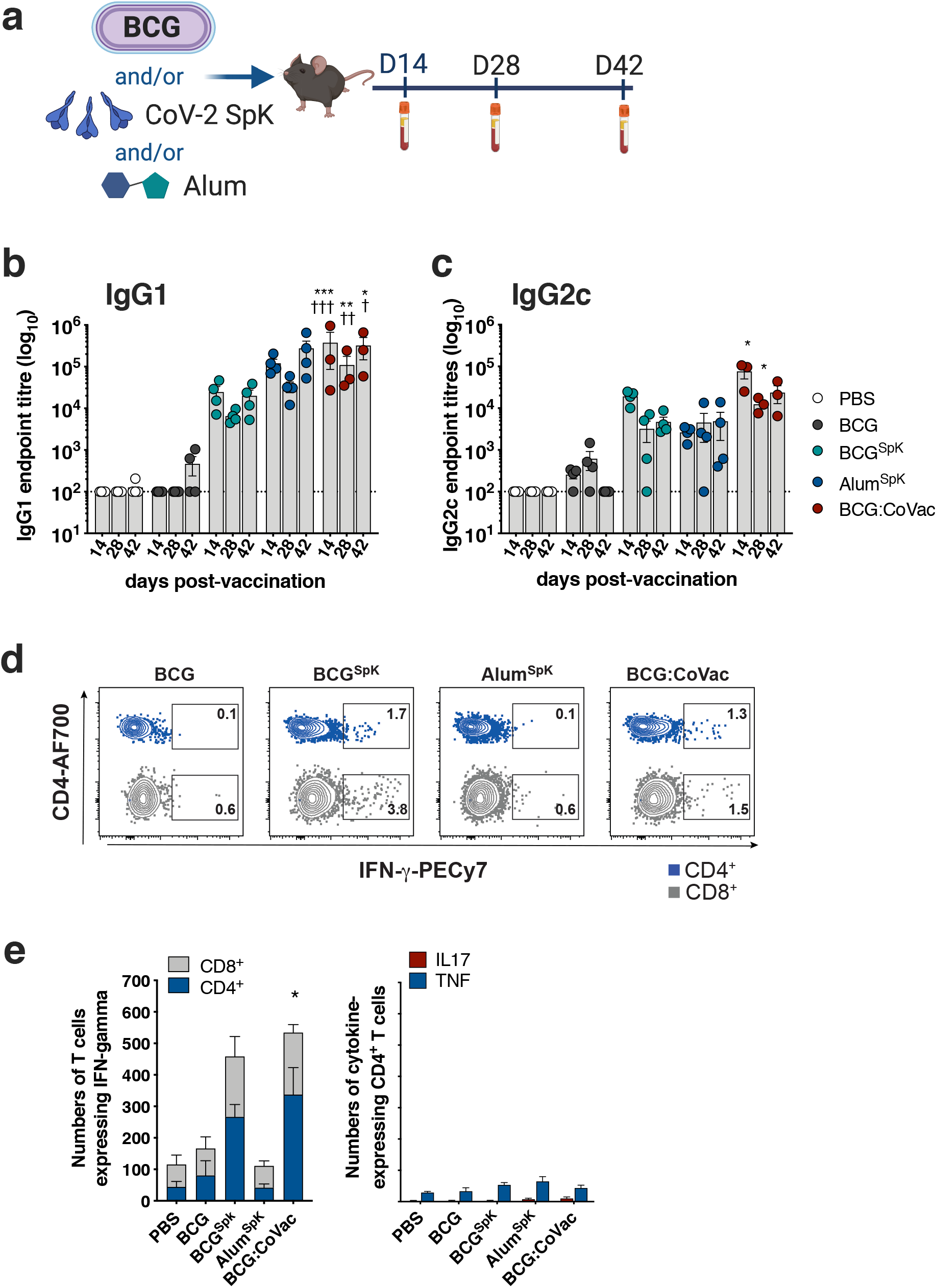
Single immunization with BCG:CoVac vaccine induces rapid development of anti-SARS-CoV-2 spike antibodies and IFN-γ-secreting T cells. **a**, C57BL/6 mice were vaccinated subcutaneously with PBS, BCG, BCG^SpK^, Alum^SpK^ or BCG:CoVac and whole blood collected at day 14, 28 and 42. **b**,**c**, Spike-specific IgG1 and IgG2c titres in plasma were determined by ELISA and estimated by the sigmoidal curve of each sample interpolated with the threshold of the negative sample ± 3 standard deviations. The dotted line shows the limit of detection. **d**, At 14 days post-vaccination PBMCs were restimulated *ex vivo* with 5 μg/mL of SARS-CoV-2 spike and cytokine production determined by flow cytometry. Representative dot plots of CD44^+^ CD4^+^ T cells and CD44^+^ CD8^+^ T cells expressing IFN-γ. **e**, Numbers of circulating CD4^+^ and CD8^+^ T cells expressing IFN-γ or CD4^+^ T cells expressing IL-17 or TNF. Data presented as mean ± s.d. Significant differences between groups compared to BCG^Spk^ *p<0.05, **p<0.01, ***p<0.01 or Alum^Spk^ †p<0.05, ††p<0.01, †††p<0.001 were determined using one-way ANOVA.

The IgG2c Ab isotype correlates with Th1-like immunity in C57BL/6 mice^13^, and such responses are considered necessary for effective protection against SARS-CoV-2 infection^14^. We therefore examined the frequency of IFN-γ-expressing T cells after a single dose of BCG:CoVac at 2 weeks post-vaccination. BCG^SpK^ and BCG:CoVac induced the generation of SpK-specific CD4^+^ and CD8^+^ T cells secreting IFN-γ (Fig. 1d, 1e), consistent with Th1 immunity observed after BCG vaccination^15^. The greatest response was observed after vaccination with BCG:CoVac, with the numbers of IFN-γ-secreting T cells significantly increased compared to vaccination with either BCG or Alum^SpK^. Low levels of the inflammatory cytokines IL-17 and TNF were observed after BCG:CoVac vaccination (Fig. 1e).

We further dissected vaccine-induced immunity by defining the cellular composition in draining lymph nodes 7 days after vaccination. Both Alum^SpK^ and BCG:CoVac induced appreciable expansion of SpK-specific germinal centre (GC) B cells (CD19^+^MHCII^+^GL7^+^CD38^-^; Fig. 2a) and plasma B cells (CD19^+^MHCII^+^CD138^+^; Fig. 2b). Cells with a T follicular helper cell (Tfh) phenotype (CD4^+^CXCR5^+^BCL6^+^) were apparent after vaccination with Alum^SpK^ or BCG:CoVac, with Tfh frequency greatest in the latter group (Fig. 2c). The total numbers of GC B cells (Fig. 2d), plasma B cells (Fig. 2e) and Tfh cells (Fig. 2f) were all significantly increased in BCG:CoVac-vaccinated mice compared to immunization with Alum^SpK^.

**Fig. 2.**
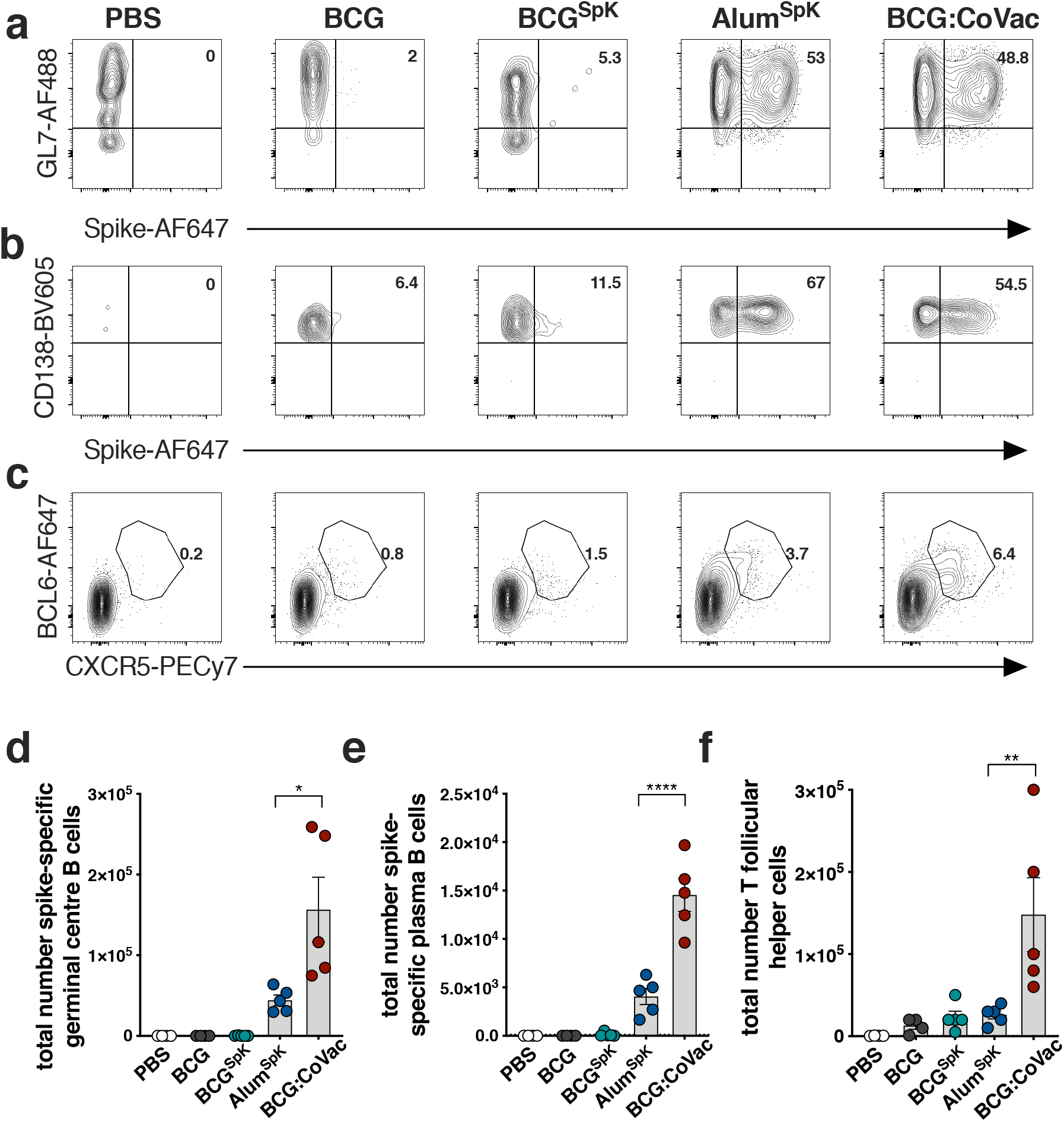
BCG:CoVac vaccination promotes expansion of T follicular helper cells and spike-specific B cells in mice. **a-c**, C57BL/6 mice were vaccinated subcutaneously with PBS, BCG, BCG^SpK^, Alum^SpK^ or BCG:CoVac and 7 days after immunization B and T cell response assessed by multicolour flow cytometry in the draining lymph node. Shown are representative dot plots of spike-specific germinal centre B cells (**a**, CD19^+^MHCII^+^GL7^+^CD38^-^), plasma B cells (**b**, CD19^+^MHCII^+^CD138^+^) and T follicular helper T cells (**c**, CXCR5^+^BCL6^+^). **d-f**, The total number of **d**, spike^+^ GC B cells, **e**, spike^+^ plasma cells and **f**, T follicular helper cells. Data presented as mean ± s.d. Significant differences between groups *p<0.05, **p<0.01, ***p<0.01 were determined by one-way ANOVA.

Overall, these data show that co-delivery of trimeric SpK antigen with BCG vaccination promotes early and pronounced anti-SARS-CoV-2 immunity, and this is further enhanced with the addition of alum.

### High-titre, SARS-CoV-2 neutralizing antibodies after a single immunization with BCG:CoVac

The elicitation of GC B cell and Tfh responses after immunization with experimental SARS-CoV-2 vaccines correlate strongly with the induction of neutralizing antibodies (NAbs)^16^. Such NAbs are a key determinant of protection induced by current vaccines used in humans^17^. We therefore measured NAb levels after a single dose of BCG:CoVac. No NAbs were detected in the plasma of mice vaccinated with BCG (Fig. 3a). Surprisingly, NAb titres were at near background levels for mice vaccinated with BCG^SpK^ (Fig. 3a), despite the high levels of IgG Ab isotypes detected in these same animals (Fig. 1). High NAb titres were detected as early as 2 weeks post-immunization upon vaccination with BCG:CoVac, and titres were significantly increased compared to vaccination with Alum^SpK^ (approximate 10-fold increase). The mean NAb titres in the plasma of BCG:CoVac-vaccinated mice were approximately 10-fold greater than those seen in SARS-CoV-2 infected humans (Fig. 3a). Although the levels of NAbs peaked at 2 weeks post-vaccination with BCG:CoVac, they remained significantly elevated up to day 42 post-immunization unlike those in the other immunized groups.

**Fig. 3.**
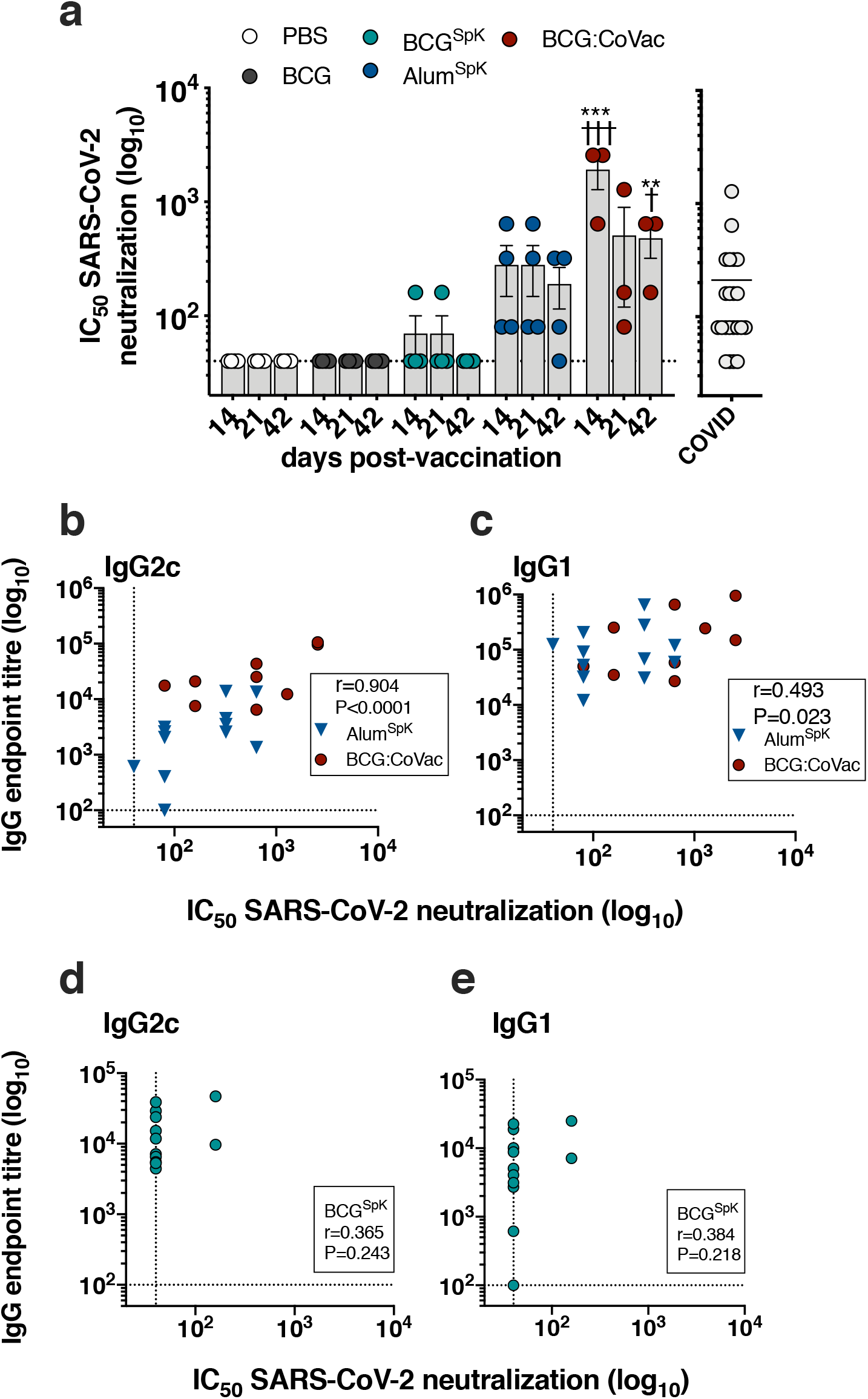
BCG:CoVac induces high titre neutralizing antibodies against live SARS-CoV-2 that correlate with the production of antigen-specific IgG2c. **a-e**, Plasma from vaccinated mice (from Fig. 1) were tested for neutralizing activity against live SARS-CoV-2 infection of VeroE6 cells. **a**, Neutralizing antibody (NAb) titres (IC_50_) were calculated as the highest dilution of plasma that still retained at least 50% inhibition of infection compared to controls. NAb titres from PCR confirmed SARS-CoV-2-infected individuals (COVID) were determined using the same method. **b**,**c**, Spearman correlations of spike-specific IgG2c or IgG1 titres and NAbs after Alum^SpK^ or BCG:CoVac vaccination. **d**,**e**, Correlation of IgG2c or IgG1 titres and NAbs after vaccination with BCG^SpK^. The dotted line shows the limit of detection. Data presented as mean ± s.d. Significant differences between groups compared to BCG^Spk^**p<0.01, ***p<0.0 or Alum^Spk^ †p<0.05, †††p<0.001 was determined by one-way ANOVA.

Since previous work suggests that the level of IgG antibody correlates with NAb titres after SARS-CoV-2 infection^18^, we examined whether a similar phenomenon was observed after vaccination with BCG:CoVac. Strong correlation (r> 0.9) was observed between IgG2c isotype and NAbs in groups vaccinated with BCG:CoVac or Alum^SpK^ (Fig. 3b), with a significant yet less robust correlation between IgG1 and NAbs for these groups (Fig. 3c). There was no correlation between NAbs and either IgG1 or IgG2c Ab for mice vaccinated with BCG^SpK^ alone (Fig. 3d, 3e).

These data suggest that alum is required for the optimal generation of NAbs after BCG:CoVac vaccination. This is a significant advantage for implementation of this vaccine candidate, due to the low cost and long standing safety record of alum^19,20^. Importantly, the potential risk of vaccine-associated enhanced respiratory disease (VAERD) caused by the selective induction of Th2 T cell responses by alum is offset by the strong Th1 immunity induced by BCG:CoVac, which in turn is driven by BCG (Fig. 1e).

### BCG:CoVac affords sterilizing immunity against SARS-CoV-2 infection in K18-hACE2 mice

Wild-type mice are not permissive to SARS-CoV-2 infection, owing to incompatibility in the receptor binding domain of the viral spike protein with the murine angiotensin-converting enzyme 2 (ACE2) ^21^. Transgenic mice expressing the human (h)ACE2 such as the K18-hACE2 mouse, are highly susceptible to SARS-CoV-2 infection, succumbing to lethal infection within 7 days post-infection^22^. We therefore assessed the protective role of BCG or BCG:CoVac vaccination in SARS-CoV-2 infection in K18-hACE2 mice. Mice were vaccinated 21 days prior to inoculation with 10^3^ PFU SARS-CoV-2 (Fig. 4a). Mice sham vaccinated with PBS succumbed to infection within 6 days with substantial deterioration in their condition with high clinical scores (Fig. 4b) and 20% weight loss (Fig. 4c). This outcome was associated with high viral titres in the airways (bronchoalveolar lavage fluid, BALF) (Fig. 4d) and lung tissues (Fig. 4e). These events led to extensive lung inflammation with substantial increases in inflammatory cells in the airways (Fig. 4f) and lung tissue (Fig. 4g), and the levels of the pro-inflammatory cytokine, IL-6, and chemokines KC (murine equivalent of IL-8) and MCP-1, in the lung tissues (Fig. 4h) and airways (Fig. S2). MCP-1 was also increased in serum (Fig. S2). These are the archetypal cytokines associated with severe human COVID-19^23^. Vaccination with BCG showed some beneficial effects and partially protected against weight loss (∼10%) and lung IL-6 and KC responses but not in other disease features. Remarkably, vaccination with BCG:CoVac 21 days prior to infection completely protected against infection, with no observable weight loss or any clinical scores throughout the duration of the experiment (Fig. 4b, 4c). These mice had no detectable virus in the airways or lungs (Fig. 4D, 4E). They had few signs of lung inflammation with moderate levels on inflammatory cells in the airways and virtually none in the lung tissue (Fig. 4g), and only baseline levels of all pro-inflammatory cytokines in the airways, lung and serum (Fig. 4h, S2). Importantly, combination of the spike protein and alum with BCG did not alter the protective efficacy of the BCG vaccine against aerosol *M. tuberculosis* in mice (Fig 4i).

**Fig. 4.**
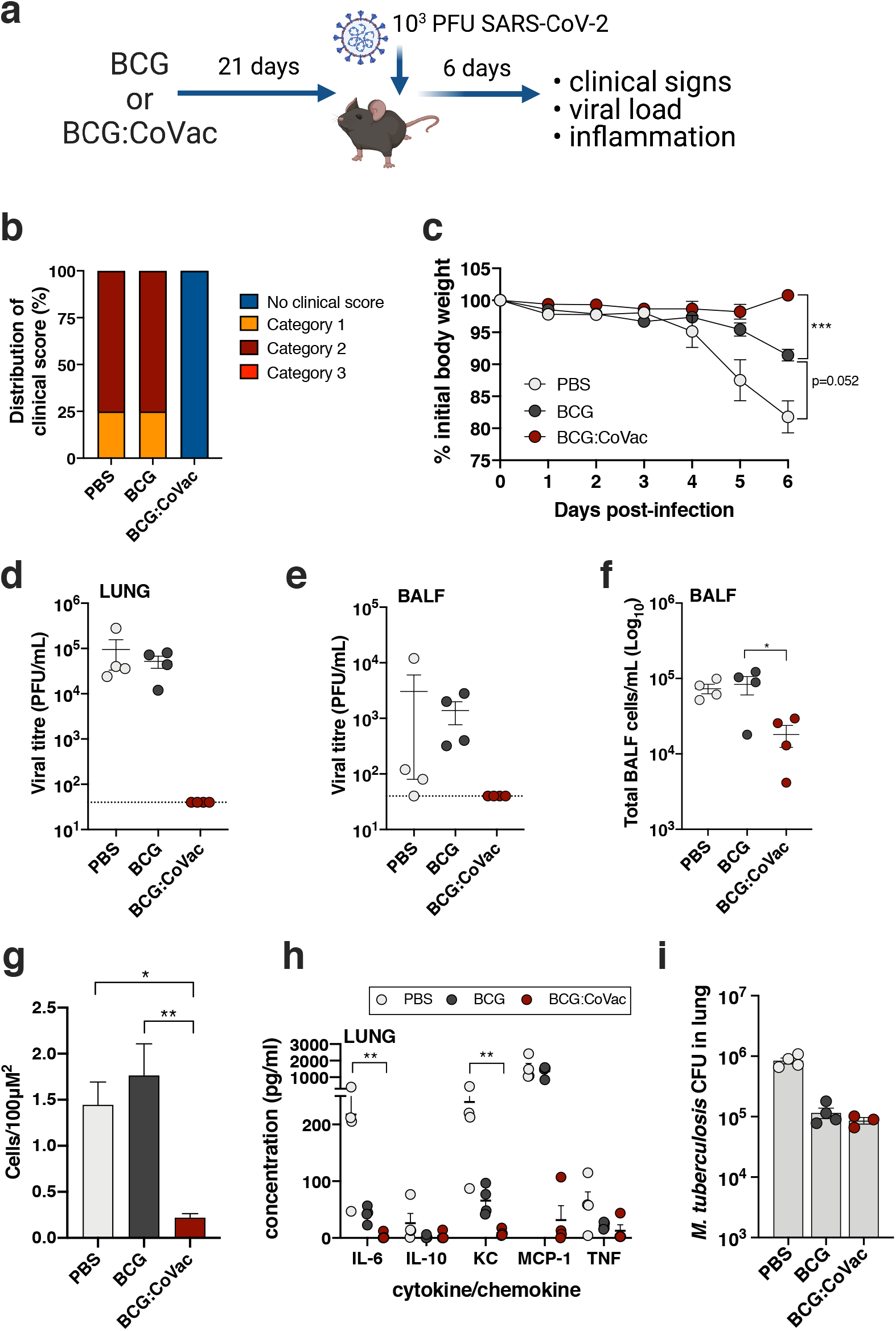
A single dose of BCG:CoVac protects against severe SARS-CoV-2 infection. **a**, Mice were immunized with sham (PBS), BCG or BCG:CoVac 21 days prior to challenge with 10^3^ PFU SARS-CoV-2. Disease outcomes were assessed 6 days later. **b**, Clinical scores at day 6 post-infection. **c**, Percentage of initial body weight loss in hemizygous male K18-hACE2 mice (n=4/group). Viral titres in lung homogenates (**d**) or bronchoalveolar lavage fluid (BALF) (**e**) were determined using plaque assay. The dotted line represents the limit of detection. **f**, Total inflammatory cells in bronchoalveolar lavage fluid (BALF). **g**, Total number of inflammatory cells in stained histological sections of lungs. **h**, Cytokine/chemokine quantification in lung homogenates. **i**, Six weeks after immunization mice were challenged with *M. tuberculosis* H37Rv by aerosol (∼100 CFU) and four weeks later the bacterial load was assessed in the lungs and presented as log_10_ of the mean CFU ± SEM. Significant differences between groups *p<0.05, **p<0.01 were determined by one-way ANOVA.

Collectively, these findings demonstrate that single dose administration of BCG:CoVac is sufficient to completely protect mice from the development of COVID-19 disease manifestations, and to neutralize infectious SARS-CoV-2 and prevent pathogenic inflammation in the lung.

### Enhancing BCG:CoVac immunity against SARS-CoV-2 by heterologous vaccine boosting

COVID-19 subunit vaccines typically display poor immunity after a single dose and require a booster to induce sufficient generation of NAbs^24^. Whilst we observed high-titre NAbs as early as two weeks post-BCG:CoVac vaccination (Fig. 3), we sought to determine if responses could be further augmented by boosting with a prototype subunit vaccine (Alum^SpK^) (Fig. 5a). At 7 days post-boost (day 28), IgG2c titres in plasma from mice primed either with BCG^SpK^ or BCG:CoVac were increased and remained elevated up to day 42 (Fig. 5b). Corresponding augmentation of NAbs was also seen in these boosted groups, with significantly elevated responses in BCG:CoVac primed mice boosted with Alum^SpK^ (Fig. 5c). Boosting Alum^SpK^ vaccination with a second dose led to a greater than 10-fold increase in NAbs in boosted mice; however, responses were significantly higher in those with the BCG:CoVac-prime, Alum^SpK-^ boost combination (Fig. 5c). Strikingly, plasma from BCG:CoVac-vaccinated mice was able to neutralize both the B.1.1.7 variant (1.3-fold decrease compared to wild-type virus) and B.1.351 variant (2.7-fold decrease) (Fig. 5d). Neutralization capacity against B.1.1.7 and B.1.351 was maintained to some extent after prime-boost with Alum^SpK^ only, however titres were approximately 10-fold less than those following the BCG:CoVac prime, Alum^SpK^ combination (Fig. 5e).

**Fig. 5.**
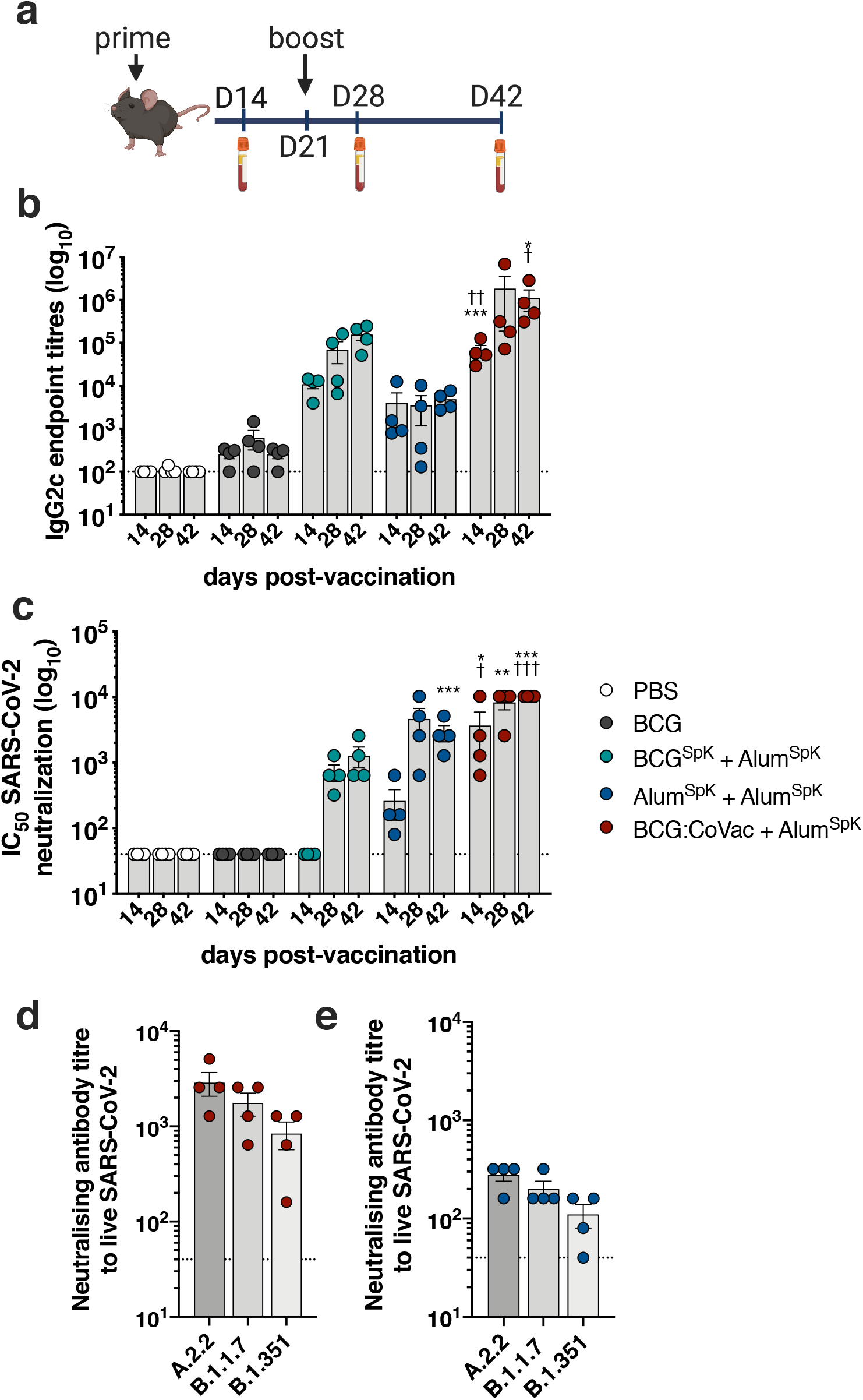
Heterologous boosting of BCG:CoVac-primed mice results in augmented SARS-CoV-2-specific IgG2c titres and neutralizing antibodies. **a**, C57BL/6 mice were vaccinated (as in Fig. 1) and at day 21 mice were boosted with Alum^Spk^. **b**, Spike-specific IgG2c titres in plasma were determined by ELISA estimated from the sigmoidal curve of each sample interpolated with the threshold of the negative sample ± 3 standard deviations. **c**, Neutralizing antibody (NAb) titres (IC_50_) were calculated as the highest dilution of plasma for all groups that still retained at least 50% inhibition of infection compared to controls. The dotted line shows the limit of detection. **d**,**e**, NAb titres against the B.1.1.7 or B.1.351 SARS-CoV-2 variants were also determined using plasma from either **d**, Alum^SpK^ or **e**, BCG:CoVac-vaccinated mice. Data presented as mean ± s.d. Significant differences between groups compared to BCG^Spk^ *p<0.05, **p<0.01, ***p<0.01 or Alum^Spk^ †p<0.05, ††p<0.01, †††p<0.001 were determined by one-way ANOVA.

Taken together, these data indicate that the antigen-specific immunity imparted by BCG:CoVac can be further enhanced by heterologous boosting with a second SARS-CoV-2 vaccine, with this vaccination regime able to induce antibodies that can neutralize key VOCs.

## DISCUSSION

Global vaccine access and distribution to low- and middle-income countries is critical for the control of the COVID-19 pandemic. Vaccines must offer effective protective immunity yet should be cheap to manufacture and have feasible cold chain management requirements. This study describes a COVID-19 vaccine formulation, BCG:CoVac, that when delivered as a single dose induces potent SARS-CoV-2 specific immunity in mice, particularly the generation of high-titre, anti-viral neutralizing antibodies. Encouragingly, the level of immune responses observed (particularly the generation of neutralizing antibodies) is equivalent to or exceeds immunity elicited by approved COVID-19 vaccines, when these candidate vaccines were tested in the murine model^24-26^. BCG:CoVac may have the additional advantage of inducing protection against other respiratory infections for which BCG is known to induce some level of protective immunity, including future pandemic viruses^11^. In addition, the possibility that prior BCG exposure may impart protection against severe COVID-19^27^, which is currently under evaluation through numerous randomised control studies^9^, raises the possibility that a BCG-based vaccine could afford protection against SARS-CoV-2 escape mutants or new pandemic coronavirus that may emerge. Indeed our data demonstrate that BCG:CoVac can neutralize two of the key VOCs that are circulating globally, namely B.1.1.7 and B.1351.

### BCG

CoVac could also provide additional benefit in countries where BCG is part of childhood immunization programs for the control of TB, based on recent findings that repeat BCG vaccination significantly reduced rates of *M. tuberculosis* infection^28^.

An advantage of our vaccine approach is the use of alum to potentiate immune responses, particularly the generation of NAbs after vaccination. Alum is a low cost, globally accessible vaccine adjuvant with an excellent safety record in humans^19^. The relative paucity of IFN-γ-secreting T cells observed after Alum^SpK^ vaccination corresponds with that previously seen with alum-precipitated vaccines using the spike protein^29^ and is consistent with the known preferential priming of Th2-type immunity by alum-based adjuvants^30^. The ability of BCG:CoVac to induce strong Th1 immunity is due to the adjuvant effect of BCG components to induce such responses^31^. This has clear importance, as T cell responses in recovering COVID-19 patients are predominately Th1-driven^32^, expression of IFN-γ was lower in severe COVID-19 cases compared to mild ones^33^ and the development of Th2 immunity correlated with VAERD^34^.We also observed only background levels of the inflammatory cytokines IL-17 and TNF after BCG:CoVac delivery, suggesting reduced levels of potentially deleterious, circulating inflammatory cytokines. Heightened expression of IL-17 correlates with severe COVID-19 disease^35^, while neutralizing IL-17 has been suggested as a possible therapy to treat acute respiratory distress syndrome in SARS-CoV-2-infected individuals^36^. In addition, the development of VAERD is also associated with Th17 immunity^37^.

Our study contributes to defining correlates of immunity in animal models that could be applied to fast-track the development of next-generation COVID-19 vaccines. A single dose of BCG:CoVac was sufficient to clear infectious SARS-CoV-2 from the lungs of K18-hACE2 mice, with no signs of clinical disease during the infection time-course, which is otherwise lethal (Fig. 4). Our findings are in agreement with previous reports that demonstrate that these mice succumb rapidly to infection, indicating that the level of NAbs elicited by BCG:CoVac is sufficient to clear SARS-CoV-2 infection in this model^21,38^. While NAb levels are known to correlate with efficacy of COVID-9 vaccines in humans^17^, other immune parameters may play an important role. Accordingly, we observed that high NAb levels in blood corelated with strong induction of GC and plasma B cells in lymph nodes, as well as heightened levels of Tfh cells. In COVID-19 convalescent individuals, the presence of memory B cells and Tfh are closely associated with NAb activity^38,39^. Further, although BCG vaccination alone did not reduce lung viral load, mice immunized with this vaccine did show an intermediate level of both weight loss and cytokine/chemokine induction during SARS-CoV-2 infection, suggesting a possible ‘dampening’ of host inflammatory responses (Fig. 4 and Fig. S2). Such an effect was seen in a human challenge model using the live-attenuated yellow fever vaccine, where BCG reduced circulating pro-inflammatory cytokines^40^. It will be of particular interest to see the outcome of ongoing clinical trials to determine if BCG vaccination can reduce COVID-19 incidence and severity^9^.

In conclusion, we describe a COVID-19 vaccine strategy based on the existing BCG vaccine, that would be broadly applicable for all populations susceptible to SARS-CoV-2 infection. Of particular note, this strategy could be readily incorporated into current vaccine schedules in countries where BCG is currently used. Further assessment in humans will determine if BCG:CoVac can impart protective immunity against not only SARS-CoV-2, but also other respiratory infections for which BCG has known efficacy.

## METHODS

### Bacterial culture

*M. bovis* BCG (strain Pasteur) was grown at 37°C in Middlebrook 7H9 media (Becton Dickinson, BD, New Jersey, USA) supplemented with 0.5% glycerol, 0.02% Tyloxapol, and 10% albumin-dextrose-catalase (ADC) or on solid Middlebrook 7H11 media (BD) supplemented with oleic acid–ADC. To prepare single cell suspensions, cultures in exponential phase (OD_600_=0.6) were washed in PBS, passaged 10 times through a 27G syringe, briefly sonicated and centrifuged at low speed for 10 min to remove residual bacterial clumps. BCG suspensions were frozen at -80° C in PBS supplemented with 20% glycerol, and colony forming units (CFU) for vaccination enumerated on supplemented Middlebrook 7H11 agar plates.

### Ethics statement

All mouse experiments were performed according to ethical guidelines as set out by the Sydney Local Health District (SLHD) Animal Ethics and Welfare Committee, which adhere to the Australian Code for the Care and Use of Animals for Scientific Purposes (2013) as set out by the National Health and Medical Research Council of Australia. SARS-CoV-2 mouse infection experiments were approved by the SLHD Institutional Biosafety Committee. COVID-19 patients were recruited through Royal Prince Alfred Hospital (RPA) Virtual, a virtual care system enabling remote monitoring of patients. The study protocol was approved by the RPA ethics committee (Human ethics number X20-0117 and 2020/ETH00770) and by the participants’ written consent. All associated procedures were performed in accordance with approved guidelines.

### Immunization

Female C57BL/6 (6-8 weeks of age) purchased from Australian BioResources (Moss Vale, Australia) or hemizygous male K18-hACE2 mice bred in-house^41^ were housed at the Centenary Institute in specific pathogen-free conditions. SARS-CoV-2 full-length spike stabilized, trimeric protein (SpK) was expressed in EXPI293F™ cells and purified as described previously^42^. Mice (n=3-4) were vaccinated subcutaneously in the footpad (s.c) with 5×10^5^CFU of BCG alone, 5 μg of SpK combined with either BCG (BCG^SpK^) or 100 μg of Alhydrogel (Alum) (Invivogen, California, USA, Alum^SpK^), or a combination of BCG (5×10^5^ CFU), SpK (5 μg) and Alyhydrogel (100 μg) (BCG:CoVac). Some mice were boosted three weeks after the first vaccination with 5 μg of SpK combined with 100 μg of Alhyhdrogel. Mice were bled fortnightly after the first immunization (collected in 10 μl of Heparin 50000 U/ml). Plasma was collected after centrifugation at 300 x *g* for 10 min and remaining blood was resuspended in 1 mL of PBS Heparin 20 U/mL, stratified on top of Histopaque 10831 (Sigma-Aldrich, Missouri, USA) and the PBMC layer collected after gradient centrifugation.

### Flow cytometry assays

Popliteal lymph nodes were collected at day 7 post immunization, and single cell suspensions were prepared by passing them through a 70 μm sieve. To assess specific B cell responses, 2×10^6^ cells were surface stained with Fixable Blue Dead Cell Stain (Life Technologies) and Spike-AF647 (1 μg), rat anti-mouse GL7-AF488 (clone GL7, 1:200, Biolegend cat#144612), rat anti-mouse MHC-II-AF700 (clone M5/114.15.2, 1:150, cat#107622), rat anti-mouse IgD-PerCP5.5 (clone 11-26c.2a, 1:200, BD, cat#564273), rat anti-mouse IgM-BV421 (clone RMM-1, 1:200, Biolegend, cat#406518), rat anti-mouse CD138-BV605 (clone 281-2, 1:200, Biolegend, cat#142516), rat anti-mouse CD19-BV785 (clone 1D3, 1:200, BD, cat#563333), rat anti-mouse CD38-APCy7 (clone 90, 1:200, Biolegend, cat#102728). To assess T cell responses, 2×10^6^ lymph node cells were stained with the following monoclonal antibodies: rat anti-mouse CXCR5-biotin (clone 2G8, 1:100, BD, cat#551960), streptavidin PECy7, rat anti-mouse CD4-AF700 (clone RM4-5, 1:200, BD cat#557956), rat anti-mouse CD8-APCy7 (clone 53-6.7, 1:200, BD cat#557654), rat anti-mouse CD44-BV605 (clone IM7, 1:300, BD cat#563058). Cells were then fixed and permeabilized using the eBioscience fixation/permeabilization kit (ThermoFischer) according to the manufacturer’s protocol and intracellular staining was performed using anti-BCL-6-AF647 (clone K112-91, 1:100, BD, cat#561525).

To assess SpK-specific cytokine induction by T cells, murine PBMCs were stimulated for 4 hrs with SpK (5 μg/mL) and then supplemented with Protein Transport Inhibitor cocktail (Life Technologies, California, USA) for a further 10-12 hrs. Cells were surface stained with Fixable Blue Dead Cell Stain (Life Technologies) and the marker-specific fluorochrome-labeled antibodies rat anti-mouse CD4-AF700 (clone RM4-5, 1:200, BD cat#557956), rat anti-mouse CD8-APCy7 (clone 53-6.7, 1:200, BD cat#557654), rat anti-mouse CD44-FITC (clone IM7, 1:300, BD cat#561859). Cells were then fixed and permeabilized using the BD Cytofix/Cytoperm™ kit according to the manufacturer’s protocol. Intracellular staining was performed using rat anti-mouse IFN-γ-PECy7 (clone XMG1-2, 1:300, BD cat#557649), rat anti-mouse IL-2-PE (clone JES6-5H4, 1:200, BD cat#554428), rat anti-mouse IL-17-PB (clone TC11-18H10.1, 1:200, cat#506918, BioLegend California, USA), rat anti-mouse TNF-PErCPCy5.5 (clone MP6-XT22, 1:200, BD cat#560659). All samples were acquired on a BD LSR-Fortessa (BD) or a BD-LSRII and assessed using FlowJo™ analysis software v10.6 (Treestar, USA).

### Antibody ELISA

Microtitration plates (Corning, New York, USA) were incubated overnight with 1 µg/mL SpK at room temperature (RT), blocked with 3% BSA and serially diluted plasma samples were added for 1 hour at 37°C. Plates were washed and biotinylated polyclonal goat anti-mouse IgG1 (1:50,000, abcam Cambridge, UK, cat#ab97238), polyclonal goat anti-mouse IgG2c (1:10,000, Abcam, cat# ab97253), or polyclonal goat anti-mouse IgG (1:350,000, clone abcam cat#ab6788) added for 1 hour at RT. After incubation with streptavidin-HRP (1:30,000, abcam, cat#405210) for 30 min at RT, binding was visualized by addition of tetramethyl benzene (Sigma-Aldrich). The reaction was stopped with the addition of 2N H_2_SO_4_ and absorbances were measured at 450 nm using a M1000 pro plate reader (Tecan, Männedorf, Switzerland). End point titres were calculated as the dilution of the sample that reached the average of the control serum ± 3 standard deviations.

### High content live SARS-CoV-2 neutralization assay

High-content fluorescence microscopy was used to assess the ability of sera/plasma to inhibit SARS-CoV-2 infection and the resulting cytopathic effect in live permissive cells (VeroE6). Sera were serially diluted and mixed in duplicate with an equal volume of 1.5×10^3^ TCID_50_/mL virus solution (B.1.319) or 1.25×10^4^ TCID_50_/mL virus solution (A2.2, B.1.1.7, B.1.351). After 1 hour of virus-serum coincubation at 37°C, 40 μL were added to equal volume of freshly-trypsinised VeroE6 cells in 384-well plates (5×10^3^/well). After 72 hrs, cells were stained with NucBlue (Invitrogen, USA) and the entire well surface was imaged with InCell Analyzer 2500 (Cytiva). Nuclei counts were obtained for each well with InCarta software (Cytiva), as proxy for cell death and cytopathic effect resulting from viral infection. Counts were compared between convalescent sera, mock controls (defined as 100% neutralization), and infected controls (defined as 0% neutralization) using the formula; % viral neutralization = (D-(1-Q))x100/D, where Q = nuclei count of sample normalized to mock controls, and D = 1-Q for average of infection controls. The cut-off for determining the neutralization endpoint titre of diluted serum samples was set to ≥50% neutralization.

### SARS-CoV-2 challenge experiments

Male hemizygous K18-hACE2 mice were transported to the PC3 facility in the Centenary Institute for SARS-CoV-2 infection. Mice were anaesthetised with isoflurane followed by intranasal challenge with 10^3^ PFU SARS-CoV-2 (VIC01/2020) in a 30 µL volume. Following infection, mice were housed in the IsoCage N biocontainment system (Tecniplast, Italy) and were given access to standard rodent chow and water *ad libitum*. Mice were weighed and monitored daily, with increased frequency of monitoring when mice developed symptoms. At day 6 post-infection, mice were euthanised with intraperitoneal overdose of pentobarbitone (Virbac, Australia). Blood was collected *via* heart bleed, allowed to coagulate at RT and centrifuged (10,000 *g*, 10 min) to collect serum. Multi-lobe lungs were tied off and BALF was collected from the single lobe *via* lung lavage with 1 mL HANKS solution using a blunted 19-gauge needle inserted into the trachea. BALF was centrifuged (300 *g*, 4°C, 7 min), and supernatants collected and snap frozen. Cell pellets were treated with 200 µL Red Blood Cell Lysis Buffer (ThermoFisher, USA) for 5 min, followed by addition of 700 µL HANKS solution to inactivate the reaction and then centrifuged again. Cell pellets were resuspended in 160 µL HANKS solution and enumerated using a haemocytometer (Sigma-Aldrich, USA). Multi-lobe lungs were collected and cut into equal thirds, before snap freezing on dry ice. Lung homogenates were prepared fresh, with multi-lobe lungs placed into a gentleMACS C-tube (Miltenyi Biotec, Australia) containing 2 mL HANKS solution. Tissue was homogenised using a gentleMACS tissue homogeniser, after which homogenates were centrifuged (300 *g*, 7 min) to pellet cells, followed by collection of supernatants for plaque assays and cytokine/chemokine measurements. The single lobe lung was perfused with 0.9% NaCl solution *via* the heart, followed by inflation with 0.5 mL 10% neutral buffered formalin through the trachea, and placed into a tube containing 10% neutral buffered formalin. Following fixation for at least 2 weeks, single lobes were transported to a PC2 facility where they were paraffin-embedded, sections cut to 3 µm thickness using a Leica microtome (Leica, Germany) and then stained using Quick Dip Stain Kit (Modified Giemsa Stain) protocol as per manufacturer’s instructions (POCD Scientific, Australia). Inflammatory cells in single lobe lungs were counted using a Zeiss Axio Imager.Z2 microscope with a 40X objective (Zeiss, Germany).

### Plaque assays

VeroE6 cells (CellBank Australia, Australia) were grown in Dulbecco’s Modified Eagles Medium (Gibco, USA) supplemented with 10% heat-inactivated foetal bovine serum (Sigma-Aldrich, USA) at 37°C/5% CO_2_. For plaque assays, cells were placed into a 24-well plate at 1.5×10^5^ cells/well and allowed to adhere overnight. The following day, virus-containing samples were serially diluted in Modified Eagles Medium (MEM), cell culture supernatants removed from the VeroE6 cells and 250 µL of virus-containing samples was added to cell monolayers. Plates were incubated and gently rocked every 15 min to facilitate viral adhesion. After 1 hr, 250 µL of 0.6% agar/MEM solution was gently overlaid onto samples and placed back into the incubator. At 72 hrs post-infection, each well was fixed with an equal volume of 8% paraformaldehyde solution (4% final solution) for 30 min at RT, followed by several washes with PBS and incubation with 0.025% crystal violet solution for 5 min at RT to reveal viral plaques.

### Cytometric bead arrays (CBAs)

CBAs were performed as per the manufacturer’s instructions (Becton Dickinson, USA). Briefly, a standard curve for each analyte was generated using a known standard supplied with each CBA Flex kit. For each sample, 10 µL was added to a well in a 96-well plate, followed by incubation with 1 µL of capture bead for each analyte (1 hr, RT, in the dark). Following capture, 1 µL of detection bead for each analyte was added to each well, followed by incubation (2 hrs, RT, in the dark). Samples were then fixed overnight in an equal volume of 8% paraformaldehyde solution (4% final solution). The following day, samples were transferred to a new 96-well plate and then transported to the PC2 facility for a second round of fixation. Samples were examined using a BD LSR Fortessa equipped with a High-Throughput Sampler (HTS) plate reader.

### *Mycobacterium tuberculosis* aerosol challenge

Eight weeks after the last vaccination mice were infected with *M. tuberculosis* H37Rv via the aerosol route using a Middlebrook airborne infection apparatus (Glas-Col, IN, USA) with an infective dose of ∼100 viable bacilli. Four weeks later, the lungs and spleen were harvested, homogenized, and plated after serial dilution on supplemented Middlebrook 7H11 agar plates. Colonies forming units (CFU) were determined 3 weeks later and expressed as log_10_ CFU.

### Statistical analysis

The significance of differences between experimental groups was evaluated by one-way analysis of variance (ANOVA), with pairwise comparison of multi-grouped data sets achieved using Tukey’s or Dunnett’s *post-hoc* test. Differences were considered statistically significant when p ≤ 0.05.

## ACKNOWLEDGMENTS

This work was supported by MRFF COVID-19 Vaccine Candidate Research Grant 2007221 (JAT, CC, PMH, SGT, MCG, ALF, WJB). We thank Florian Krammer of the Icahn School of Medicine, Mt Sinai for provision of the pCAGGS vector containing the SARS-CoV-2 Wuhan-Hu-1 Spike Glycoprotein Gene. We acknowledge the support of the University of Sydney Advanced Cytometry Facility and the animal facility at the Centenary Institute. We thank Sunil David (ViroVax LLC) and Wolfgang Leitner (NIAID, NIH) for helpful discussions. PMH is funded by a Fellowship and grants from the National Health and Medical Research Council (NHMRC) of Australia (1175134) and by UTS. JAT, BMS and WJB are supported by the NHMRC Centre of Research Excellence in Tuberculosis Control (1153493). The establishment of the PC3 COVID-19 facility was supported by UTS and the Rainbow Foundation.

## AUTHOR CONTRIBUTIONS

CC, MDJ, AOS, SGT, PHM and JAT designed the study. CC, MDJ, AOS, DHN, ALF, AA, RS, NDB, AG, KP, BMS, JKKL, SGT performed the experiments. All authors contributed to data analysis/interpretation. CC, MDJ, PHM and JAT wrote the first manuscript draft and all authors provided revision to the scientific content of the final manuscript.

## FIGURE LEGENDS

**Fig. S1.**
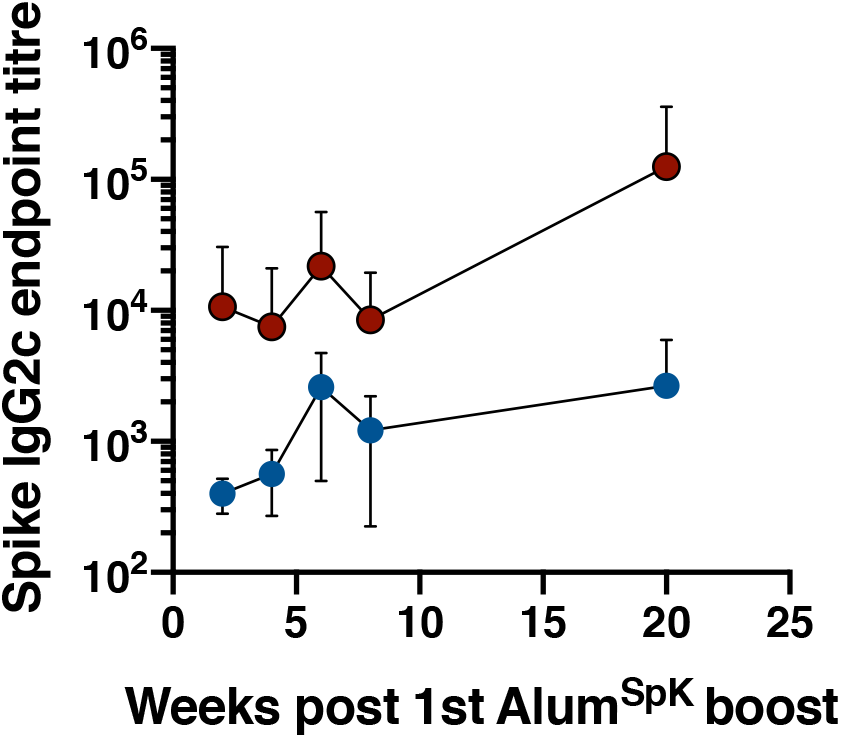
BCG promotes spike-specifica antibody reponses. Mice were vaccinated s.c with BCG (5×10^5^ CFU) and 12 week later were vaccinated twice, 3 weeks apart with s.c with SARS-CoV-2 spike protein (5 μg) formulated in alum (100 μg; Alum^SpK^). At the indicated timepoints the titre of spike-specific IgG2c in sera was determined by ELISA.

**Fig. S2.**
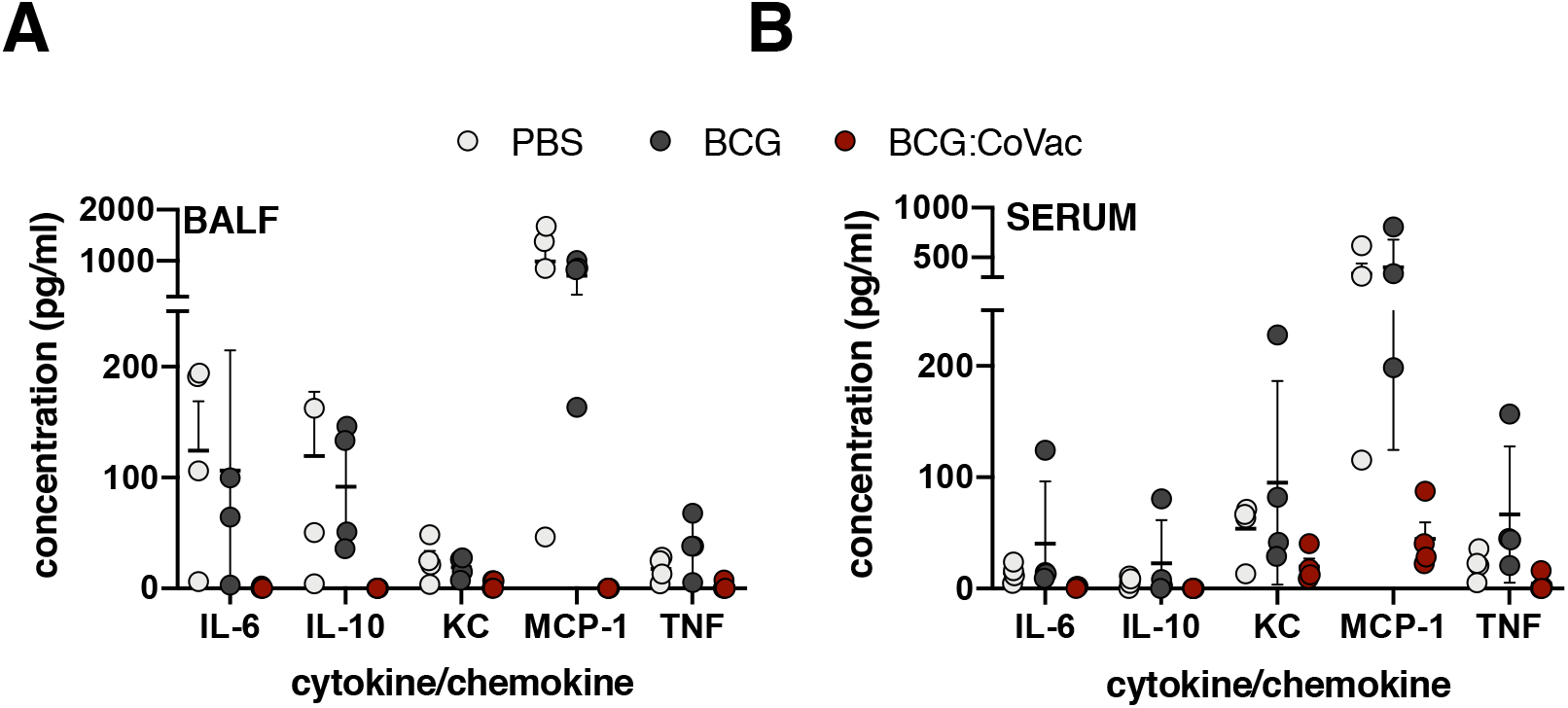
Single dose BCG:CoVac prevents the development of clinical disease in SARS-CoV-2 infected K18-hACE2 mice. **a**, Mice were immunised with sham (PBS), BCG or BCG:CoVac 21 days prior to challenge with 10^3^ PFU SARS-CoV-2. Cytokine/chemokine levels were determined in BALF (**a**) and serum (**b**) by cytometric bead array.

## Notes

### Competing Interest Statement

The authors have declared no competing interest.

